# Computer modeling of whole-cell voltage-clamp analyses to delineate guidelines for good practice of manual and automated patch-clamp

**DOI:** 10.1101/2020.04.27.062182

**Authors:** Jérôme Montnach, Maxime Lorenzini, Adrien Lesage, Isabelle Simon, Sébastien Nicolas, Eléonore Moreau, Céline Marionneau, Isabelle Baró, Michel De Waard, Gildas Loussouarn

## Abstract

The patch-clamp technique has contributed to major advances in the characterization of ion channels. The recent development of high throughput patch-clamp provides a new momentum to the field. However, whole-cell voltage-clamp technique presents certain limits that need to be considered for robust data generation. One major caveat is that current amplitude profoundly impacts the precision of the analyzed characteristics of the ion current under study. For voltagegated channels, the higher the current amplitude is, the less precise the characteristics of voltage-dependence are. Similarly, in ion channel pharmacology, the characteristics of dose-response curves are hindered by high current amplitudes. In addition, the recent development of high throughput patch-clamp technique is often associated with the generation of stable cell lines demonstrating high current amplitudes. It is therefore critical to set the limits for current amplitude recordings to avoid inaccuracy in the characterization of channel properties or drug actions, such limits being different from one channel to another. In the present study, we use kinetic models of a voltage-gated sodium channel and a voltage-gated potassium channel to edict simple guidelines for good practice of whole-cell voltage-clamp recordings.

## INTRODUCTION

The patch-clamp technique has contributed to major advances in the characterization of ion channel biophysical properties and pharmacology, thanks to the versatility of the readouts: (i) unitary currents allowing the study of a single channel conductance, open probability and kinetics, and (ii) whole-cell currents allowing characterization of a population of channels, macroscopic properties of the gates, or pharmacology (Hille, 2001).

As for any techniques, some practical limits have to be taken into account. As schematized in Figure 1, a major caveat in the whole-cell configuration of the voltage-clamp technique is due to the fact that the pipette tip in manual patch-clamp, or the glass perforation in planar automated patch-clamp, creates a series resistance (R_S_) in the order of the MΩ. Consequently, according to the Ohm’s law, when a current flowing through the pipette is in the order of the nA, it leads to a voltage drop of several mV at the pipette tip or the glass perforation. The actual voltage applied to the cell membrane (V_m_) is therefore different than the voltage clamped by the amplifier and applied between the two electrodes (pipette and bath electrodes, V_cmd_).

**Figure 1:**
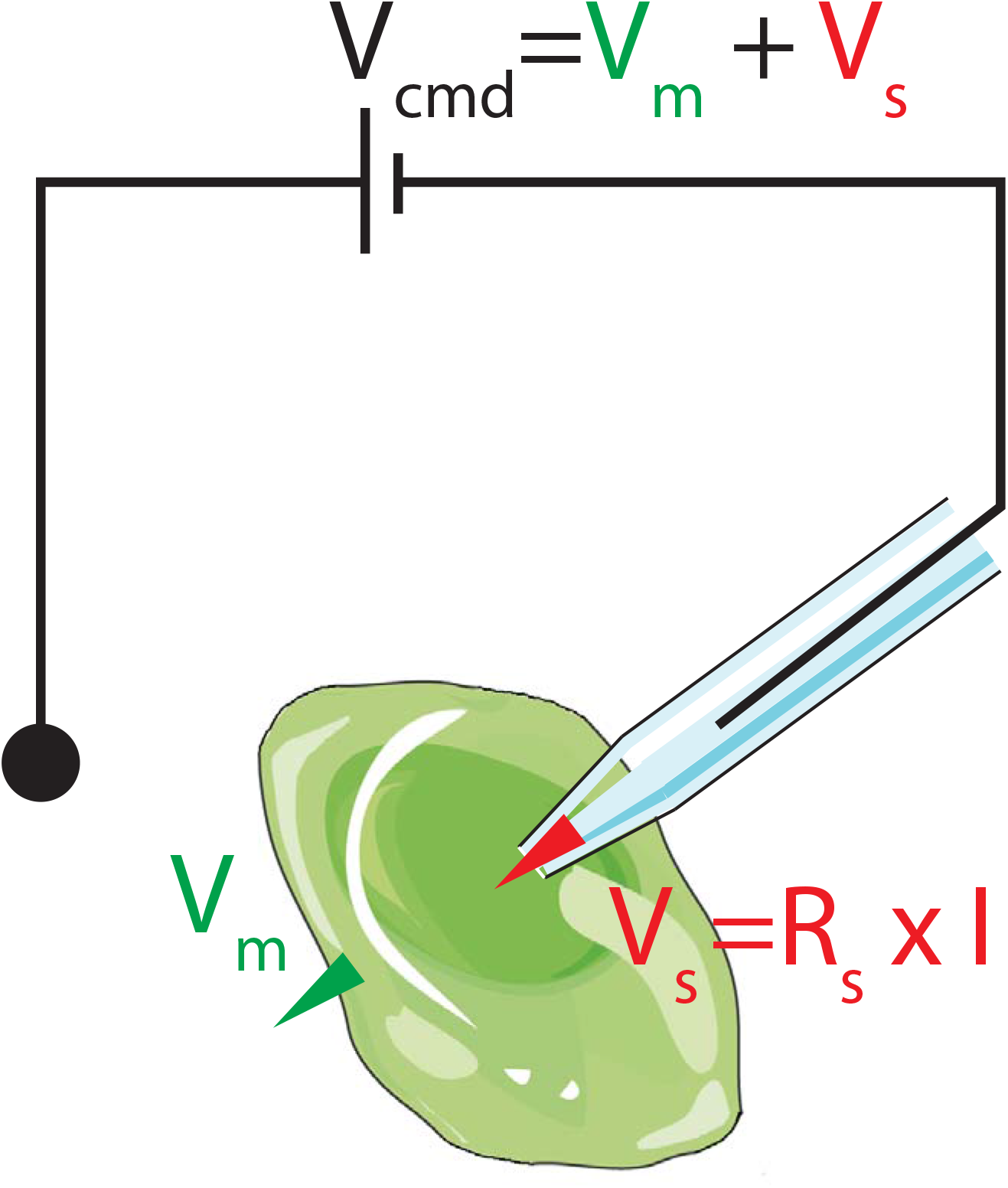
Simplified scheme of a cell studied in voltage-clamp, illustrating how the potential clamped between the two electrodes is split between the membrane potential and a potential generated at the pipette tip.

This caveat was described early on when the patch-clamp technique was developed (Ebihara and Johnson, 1980). However, with the development of automated patch-clamp for the industry and academia, the technique that was formerly reserved to specialized electrophysiologists, has been popularized to scientists that are not always aware of the limits of the technique. In that respect, we witness the multiplication of publications that report ionic currents in the range of several nA, that undoubtedly have led to incorrect voltage clamp and erroneous conclusions. Early on, this problem was partially solved by the development of amplifiers having the capacity to add a potential equivalent to the one which is lost (V_S_), a function which is called R_S_ compensation (Sherman et al., 1999). However, compensation rarely reaches 100% and high-throughput systems have limited compensation capabilities due to the fact that there is only one amplifier for an array of cells.

Here, we used a mathematical model to study in detail the impact of various levels of R_S_ and current amplitude on the steady-state activation and dose-response curves of the cardiac voltage-dependent sodium current I_Na_, as well as the steady-state activation curve of the cardiac voltage-dependent potassium current I_to_. We then predicted the impact of various levels of R_S_ on the current conducted by transiently expressed Na_v_1.5 channels and compared this prediction to whole-cell voltage-clamp recordings obtained in manual patch-clamp analyses. Finally, we looked at the impact of R_S_ in whole-cell voltage-clamp recordings of Na_v_1.5 currents obtained in automated patch-clamp. This study highlights potential incorrect interpretation of the data and allows proposing simple guidelines for the study of voltage-gated channels in patch-clamp, which will help in the design of experiments and rationalize data analyses to avoid misinterpretation.

## RESULTS

The aim of this study was to use kinetic models of specific ion currents to generate current ‘recordings’ that take into account the voltage error made using the whole-cell configuration of the patch-clamp technique. We then used and compared these data and experimental observations to propose simple rules for good quality patch-clamp recordings.

In order to calculate the current (I) recorded at a given voltage, in a cell, we used Hodgkin-Huxley models of voltage-gated channels (O’Hara et al., 2011). For this calculation, we need to determine the actual membrane potential (V_m_) but we only know the potential applied between the pipette and reference electrode (V_cmd_), illustrated in Figure 1. The voltage error between V_m_ and V_cmd_ is the voltage drop at the pipette tip (V_S_), due to the series resistance (R_S_). So, I is the resultant of both the membrane resistance variations (due to channels opening or closing, R_m_) and R_S_. For voltage-gated channels, R_S_ varies with voltage and time. So we envision that I value can only be obtained through an iterative calculation at each time step (see supplemental information for further details).

We started to model the current conducted by cardiac voltage-gated sodium channels (Na_v_1.5 for the vast majority) for a combination of values of series resistance (R_S_) and various current amplitude ranges (depending on the amount of active channels in the membrane, Figure 2A). First, when R_S_ is null (the optimal condition, which can only be attained experimentally by 100% R_S_ compensation), the voltage error is null and the shapes of the recordings are identical, independent of the current amplitude (in green in Figure 2B). Consistent with voltage error being proportional to both R_S_ and current values, we observed that combined increase in R_S_ and current amplitude leads to alteration in the current traces, due to a deviation of V_m_ from V_cmd_ (Figure 2B). When R_S_ is equal to 2 MΩ (in orange), alterations in the shape of the currents are observed only when current amplitude reaches several nA (high expression of ion channels, bottom). Further increasing R_S_ to 5 MΩ, the alterations are observed already in the medium range of current, and emphasized when currents are large (middle and bottom, in red). As illustrated in Figure 2B, when I and R_S_ are elevated, the voltage applied to the membrane, V_m_, can reach −7 mV only, whereas the applied voltage command, V_cmd_, is −40 mV (bottom right, Vm inset). Thus, in these conditions, V_S_, the voltage deviation represents 33 mV, which is not at all negligible (Figure 2C).

**Figure 2:**
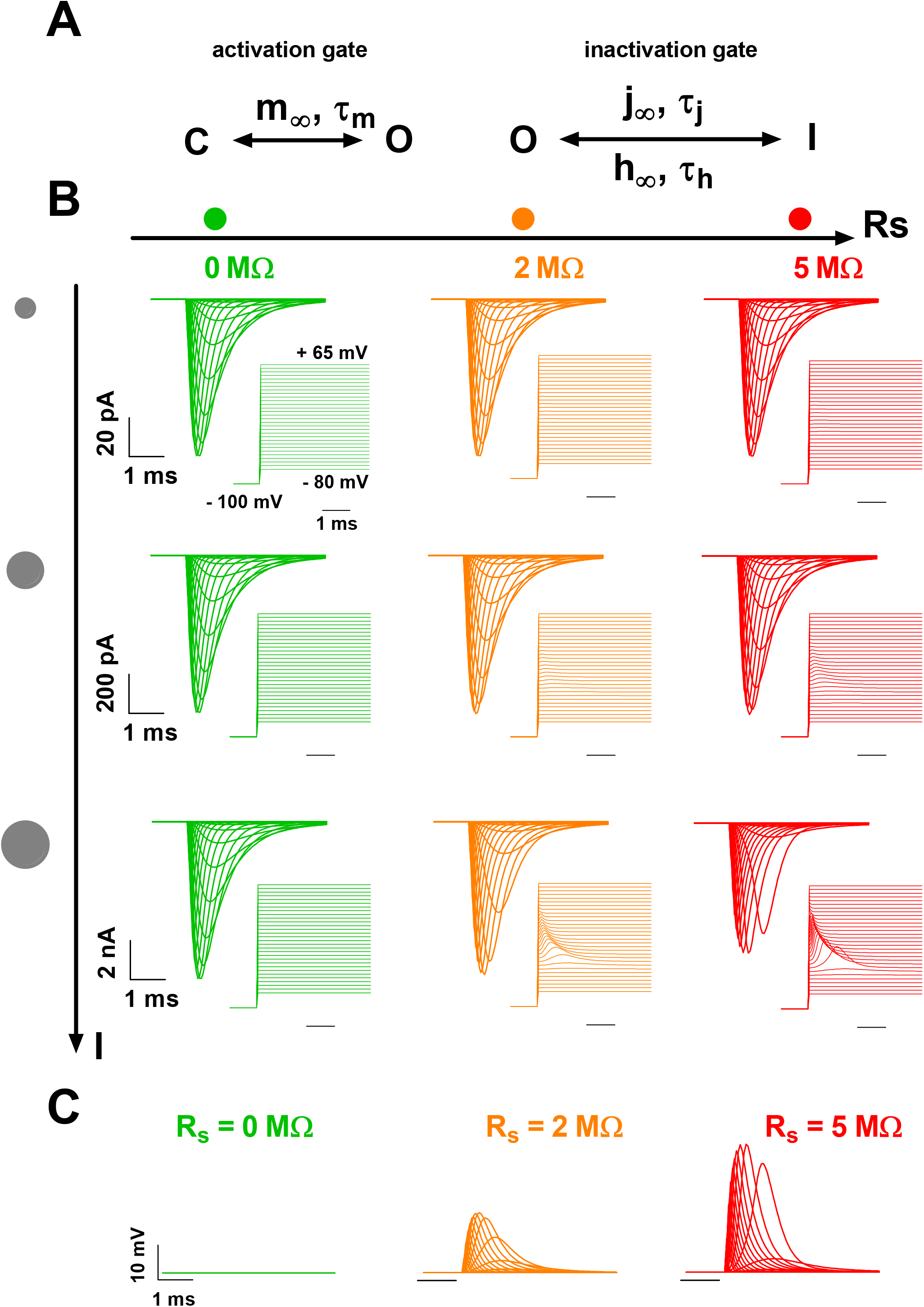
Kinetic model of cardiac I_Na_ current (Na_v_1.5) - computed effects of increasing series resistance and current amplitude range on current recordings. **A.** Scheme of the kinetic model, with independent activation and inactivation gates (C, O and I: closed, open, and inactivated states, respectively; O’Hara et al., 2011). **B.** Superimposed computed I_Na_, for increasing current amplitude range (I, see vertical scales, top to bottom) and series resistance (R_S_, left to right). The activation voltage protocol shown (in inset, holding potential: −100 mV; 50-ms pulse at the indicated potentials; one sweep every 2 s) corresponds to the potential at the membrane, not the command potential between the two electrodes. It is thus altered when (R_S_ x I) is elevated, its maximum deflection reaching 33 mV at V_cmd_ = −40 mV (cf text). **C.** Superimposed computed voltage deviation for the highest amplitude (10 nA) and various R_S_, as in B.

The impact of R_S_ on current amplitude is the highest for large amplitudes (Figure 3A, 10-nA range), e.g. at potentials between −40 and −20 mV. At such potentials, activation and inactivation time course are also altered by high R_S_ values. Altogether, this leads to major artefactual modifications of the current-voltage and activation curves (Figure 3B-C). Indeed, except when R_S_ is null, increasing the current amplitude range, from 0.1 to 10 nA, shifts the voltage-dependence of activation (V_0.5_) towards hyperpolarized potentials (Figure 3C). For the largest currents, series resistance of 2 and 5 MΩ induces a −8 mV and −16 mV shift of V_0.5_, respectively. The slope factor k is also drastically reduced by a factor of 1.7 and 3.6, respectively. As a result, for the largest current and series resistance of 5 MΩ, an all-or-nothing-type response is generated by changing the step voltage from −50 (almost no activation) to −45 mV (almost full activation).

**Figure 3:**
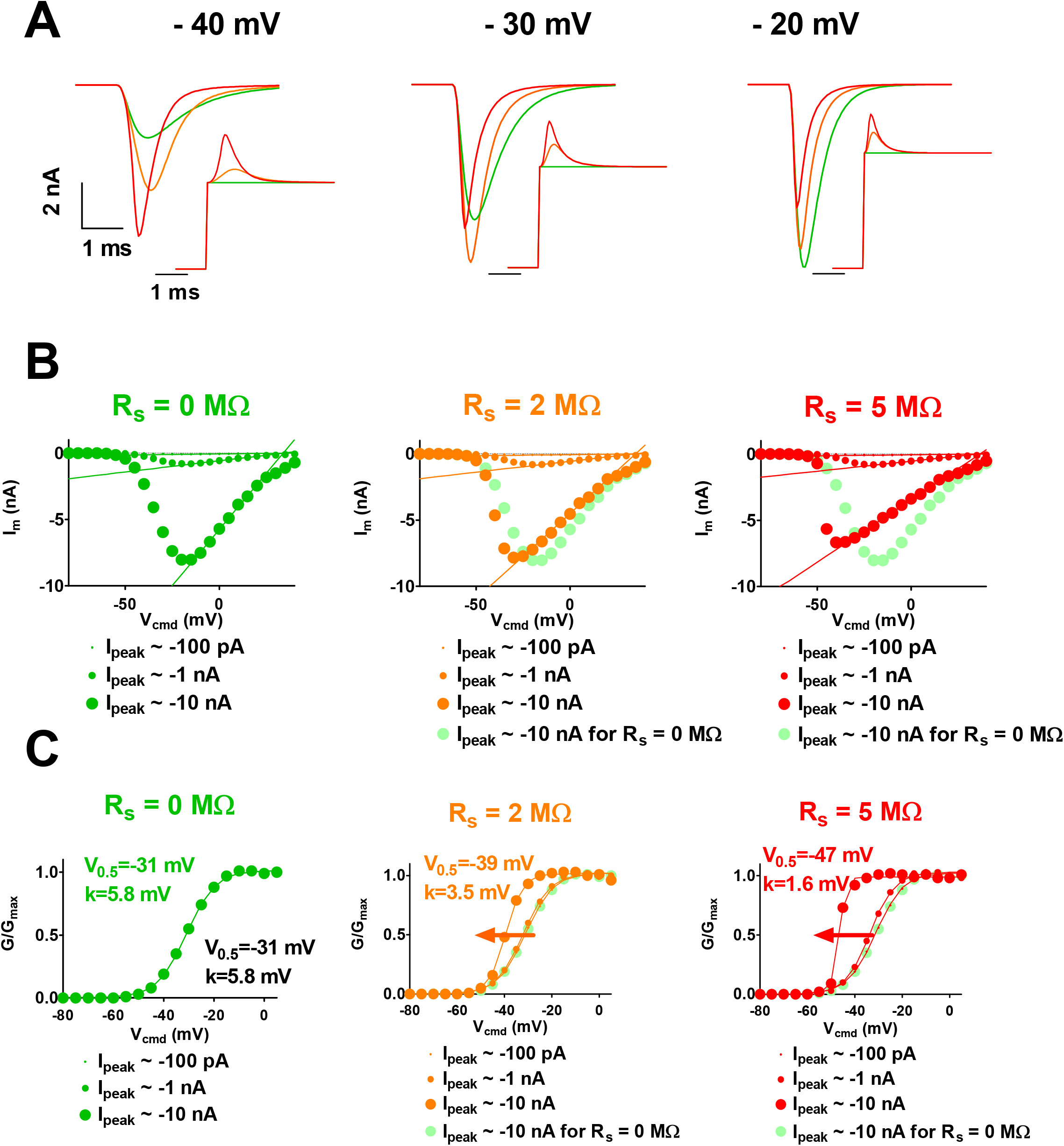
Kinetic model of cardiac I_Na_ current (Na_v_1.5) - computed effects of increasing series resistance and current amplitude range on apparent current parameters. **A.** Computed time course of I_Na_ current at the indicated command potential, for various values of R_S_, as indicated by colors. **B.** Peak current/voltage (I/V) relationships in the three conditions. Optimal I/V curve is repeated (light green), allowing easy comparison. **C**. Activation curves (G/G_max_ *vs* V_cmd_) in various conditions as indicated by color and symbol size. Theoretical half-activation potential (V_0.5_) and slope (k) values in black. V_0.5_ and k for R_S_ of 0, 2 and 5 MΩ and current range of 10 nA are indicated in colors.

Besides impact on the characteristic of voltage dependent activation, R_S_ may also impact channel pharmacological characteristics. In order to model this impact, we had to draw for various values of R_S_, the relationship between the theoretical values of the peak sodium current, I_peak_ (with no voltage error) and the measured values of I_peak_. We calculated this relationship at a potential that could be used to establish the dose-response curve, here −20 mV. First, when R_S_ is null, the voltage error is null and both values (theoretical and computed values) are the same. As R_S_ increases, the measured I_Na_ curve is inflected accordingly (Figure 4A, middle and right).

**Figure 4:**
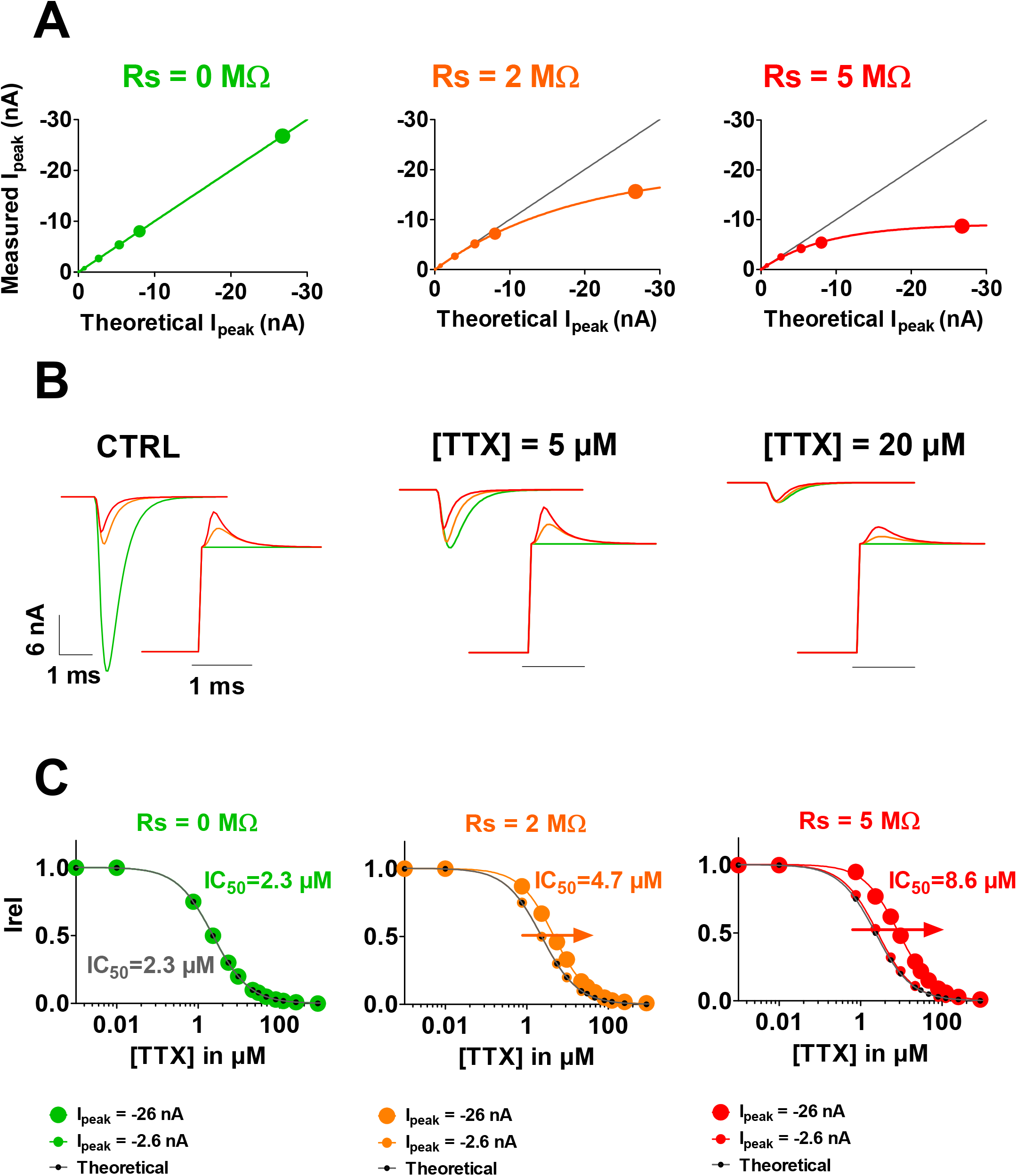
Kinetic model of cardiac I_Na_ current (Na_v_1.5) - computed effects of increasing series resistance and current amplitude on apparent TTX effects. **A.** Relationships between the theoretical values of peak I_Na_ (with no voltage error) and the measured values of I_Na_ for three different values of R_S_ (0, 2 and 5 MΩ). **B.** Computed recordings of the I_Na_ current at the indicated TTX concentrations, for various values of R_S_, as indicated by colors. **C**: Computed dose-response curves for three R_S_ conditions (R_S_ = 0, 2 and 5 MΩ) and two amplitude ranges (I peak = −2.6 and −26 nA).

We used this relationship to look at the impact of R_S_ on the apparent effects of a channel blocker, tetrodotoxin (TTX) on the sodium current. We started with published data on TTX (Wang et al., 2013) to generate theoretical (R_S_ = 0 MΩ) dose-response curve and current traces in the presence of various concentrations of TTX (Figure 4B-C, green). Then we used the relationship between the theoretical and values of I_peak_ at R_S_ of 2 and 5 MΩ (established in Figure 4A), to build the dose-response curves, for these R_S_ values. When there is no TTX (Figure 4B, left), current amplitude is high and voltage-clamp is not efficient, thus there are major differences between theoretical (green) and measured (orange and red) amplitudes. When inhibitor concentration increases, remaining current amplitudes decrease and voltage-clamp is improved. Hence, theoretical and measured values are similar, whatever the R_S_ values (Figure 4B, right). This leads to an artefactual shift of the resulting dose-response curve toward higher concentrations (Figure 4C). For I_peak_ of 26 nA, R_S_ of 2 and 5 MΩ induce an increase of IC_50_ by a factor of 2.0 and 3.9, respectively. For low I_peak_ (2.6 nA), these modifications are minimal. Since R_S_ values of 10 MΩ can be easily met in automated patch clamp, we also calculated IC_50_ for a I_peak_ of 26 nA and obtained an increase of IC_50_ by a factor of 5.3.

We applied the same modeling strategy to study the impact of R_S_ on the ‘measurement’ of the voltage-gated K^+^ current I_to_, using a Hodgkin-Huxley model of this current (Figure 5A; O’Hara et al., 2011). As for the Na^+^ current, we modeled the I_to_ current for a combination of values of series resistances and current amplitude ranges. Again, when R_S_ is null, the voltage error is null and the shapes of the recordings are identical, independent of the current amplitude (in green in Figure 5B). Consistent with voltage error being proportional to both R_S_ and current, we observed that combined increase of R_S_ and current amplitude leads to alteration in the recordings, due to a deviation of V_m_ from V_cmd_ (Figure 5B). I_to_ current characteristics, nonetheless, are less sensitive to R_S_ and current amplitude as compared to I_Na_: when R_S_ is equal to 5 MΩ (in orange), alteration in the shape of the recordings is significant only when current amplitude is 10-fold higher than Na^+^ currents (in red). When R_S_ reaches 15 MΩ and current amplitude is equal to several tens of nA, conditions routinely observed in automated patch-clamp with stable cell lines (Oliver et al., 2017; Ranjan et al., 2019), the model predicts major modification of the activation curve and apparition of a delayed inactivation (Figure 5B, bottom right). When R_S_ is not null, increasing peak current amplitude range up to 100 nA leads to major shift in voltage-dependence of activation as the following: for a peak current of 100 nA, R_S_ of 5 and 15 MΩ induces a −9 mV and −18 mV shift of the half activation potential, respectively. The slope is also drastically decreased by a factor of 1.8 and 2.5, respectively (Figure 5C). Noteworthy, when R_S_ is 15 MΩ and amplitudes are in the order of several tens of nA, major voltage deviation occurs, decreasing the current amplitude by a factor of ten. This may give the impression that the current is not high and thus voltage error negligible.

**Figure 5:**
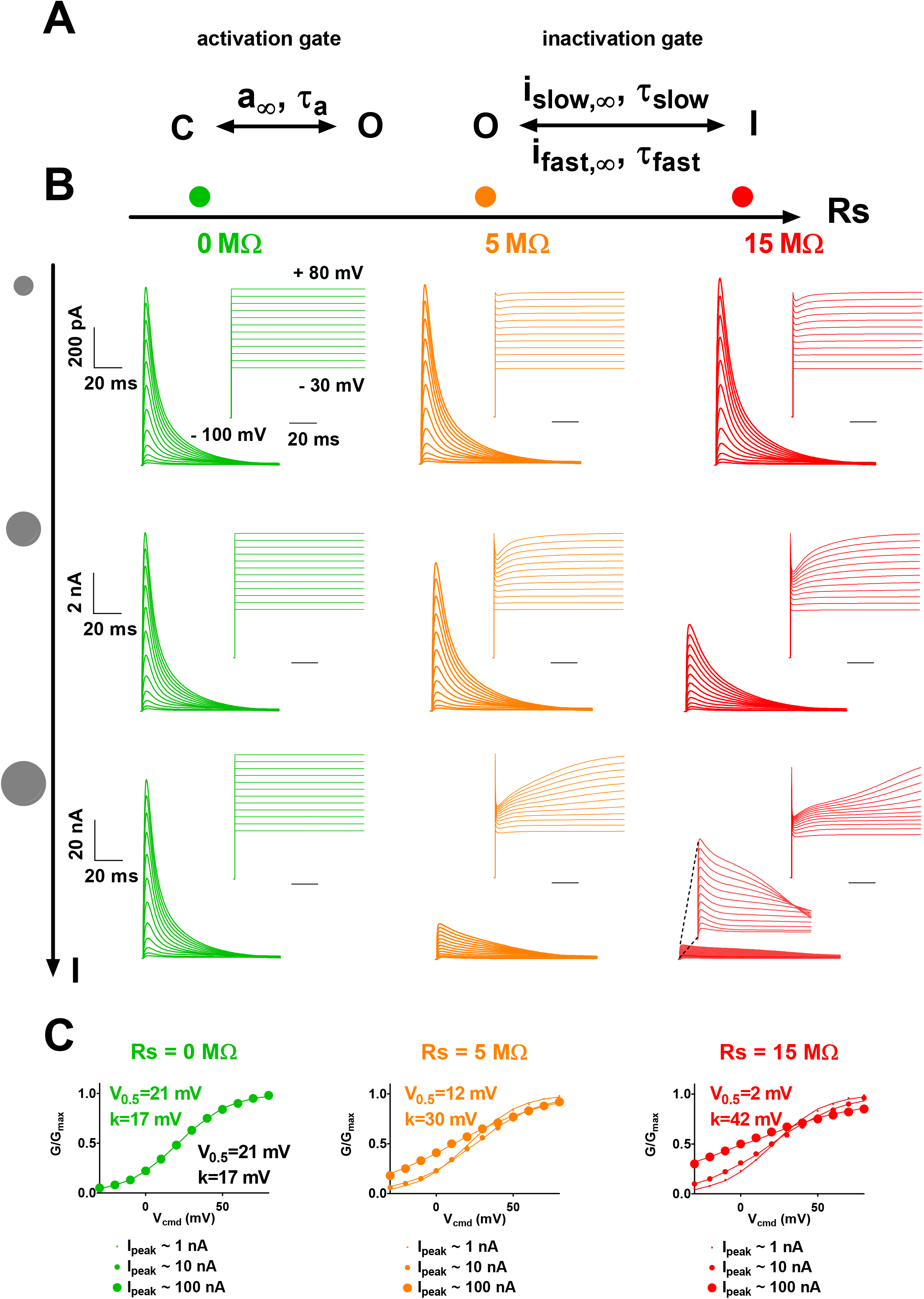
Kinetic model of cardiac I_to_ current; computed effects of increasing series resistance and current amplitude on current recordings. **A.** Scheme of the kinetic model, with independent activation and inactivation gates (O’Hara et al., 2011). **B.** Computed superimposed recordings of the I_to_ current, for various current amplitudes and series resistances (R_S_), as indicated. The activation voltage protocol shown (holding potential: −100 mV; 100-ms pulse at the indicated potentials; one sweep every 2 s) corresponds to the potential experienced by the membrane (V_m_), not the potential between the two electrodes (V_cmd_). It is thus altered when (R_S_ x I) is elevated. **C.** Activation curves (G/G_max_ *vs* V_cmd_) relationships in the three conditions represented in B. Theoretical half-activation potentials (V_0.5_) and slopes (k) are indicated in black. V_0.5_ and k for R_S_ of 0, 5 and 15 MΩ and I peak of 100 nA are indicated in colors.

In order to test whether the model reproduces experimental data, we used a set of data of heterologously-expressed Na_v_1.5 currents recorded in COS-7 cells using manual voltage-clamp (qualitatively validated or not, to include highly ‘artefacted’, erroneous data). When using transient transfection systems, recorded currents are very variable from cell to cell, with peak currents measured at −20 mV ranging from 391 pA to 17.8 nA in the chosen cell set (52 cells). We used this variability to study the effect of current amplitude on the activation curve properties. First, in order for the model to be as close as possible to experimental data, we modified the previously published Hodgkin-Huxley model, to match the properties of the Na_v_1.5 current obtained in optimal experimental conditions (Figure 6). We used as reference group, the cells presenting peak current amplitudes in a range smaller than 1 nA (7 cells), and with R_S_ compensation allowing residual R_S_ of around 2 MΩ. The initial model (Figure 3) suggests negligible alteration of V_0.5_ and k in these conditions. The model was then optimized to obtain V_0.5_, k and inactivation time constants that are similar to averaged values of the 7 reference cells (Figure 6B,C).

**Figure 6:**
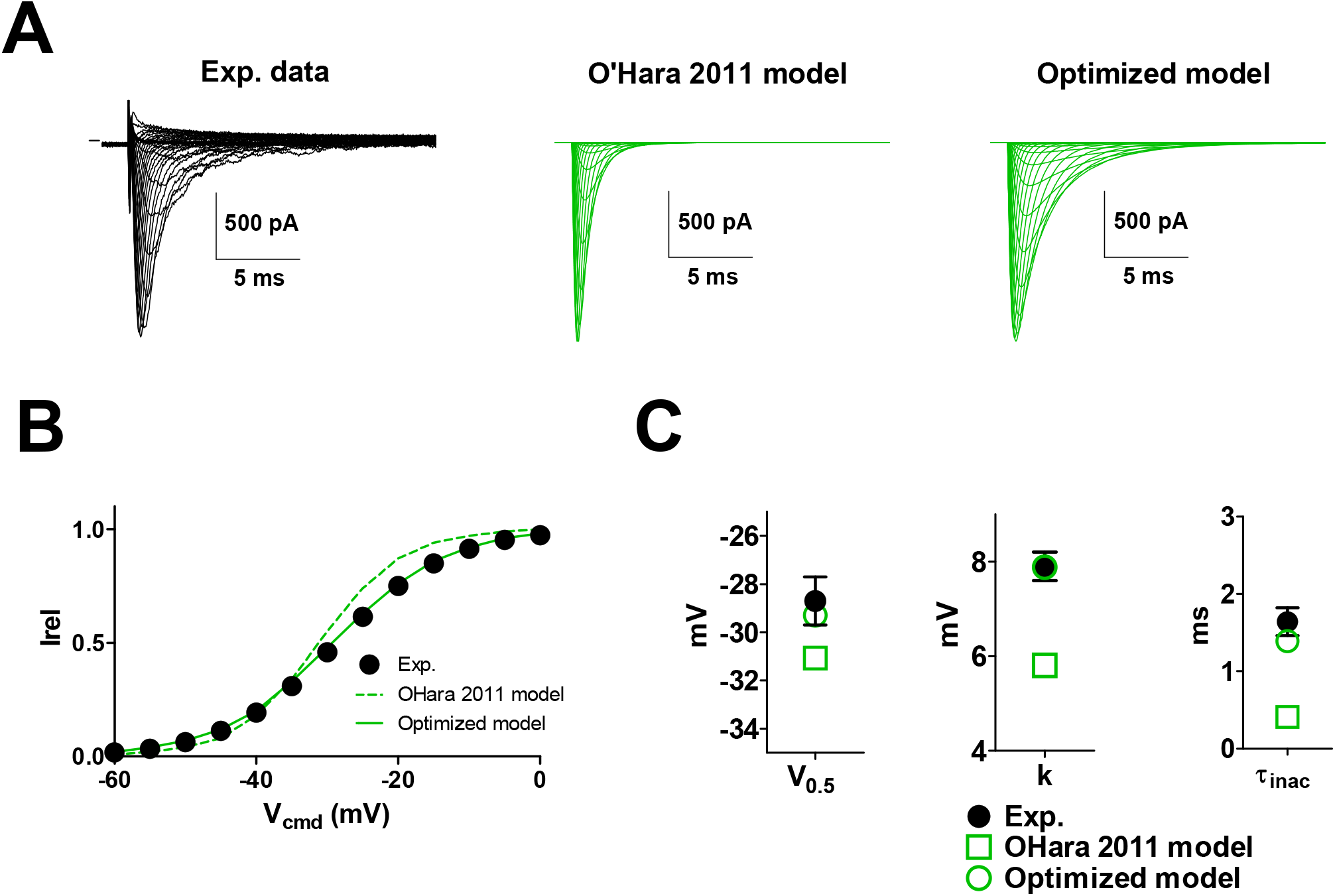
Optimization of Kinetic model presented in Figure 2 to fit the recordings of heterologously-expressed Na_v_1.5. **A**: *Left*, representative, superimposed recordings of heterologously-expressed Na_v_1.5 during an activation voltage protocol (same as in Figure 2). *Middle*, computed recordings using the same protocol, and the equation of (O’Hara et al., 2011). *Right*, computed recordings using the same protocol, and the optimized model, to fit the biophysical characteristics of heterologously-expressed Na_v_1.5. **B**: Activation curves in the three conditions presented in A. **C**. half-activation potentials, slopes and time constants of inactivation in the three conditions.

We then split the 52 cells in 6 groups according to current amplitude range (the 7 reference cells, then 4 groups of 10 cells, and a last group of 5 cells) and plotted, for each group, the mean V_0.5_ and k values as a function of mean current amplitude (Figure 7). We observed a decrease in both V_0.5_ and k when current range increases. These relationships were successfully fitted by the computer model when R_S_ was set to 2 MΩ, which is close to the experimental value, after compensation (R_S_ = 2.3 ± 0.2 MΩ). In these conditions, if we accept maximal inward peak current amplitudes up to 7 nA, error on V_0.5_ is below 10 mV and k remains greater than 5 mV. Experimentally, current amplitudes larger than 7 nA should be prevented or discarded, to prevent larger errors on the estimate of V_0.5_ and k. Nevertheless, the benefit of such a representation (Figure 7) is obvious as a correlation can be drawn between current amplitude and V_0.5_ and k values, with a more reliable estimate of these values at low current amplitude levels.

**Figure 7:**
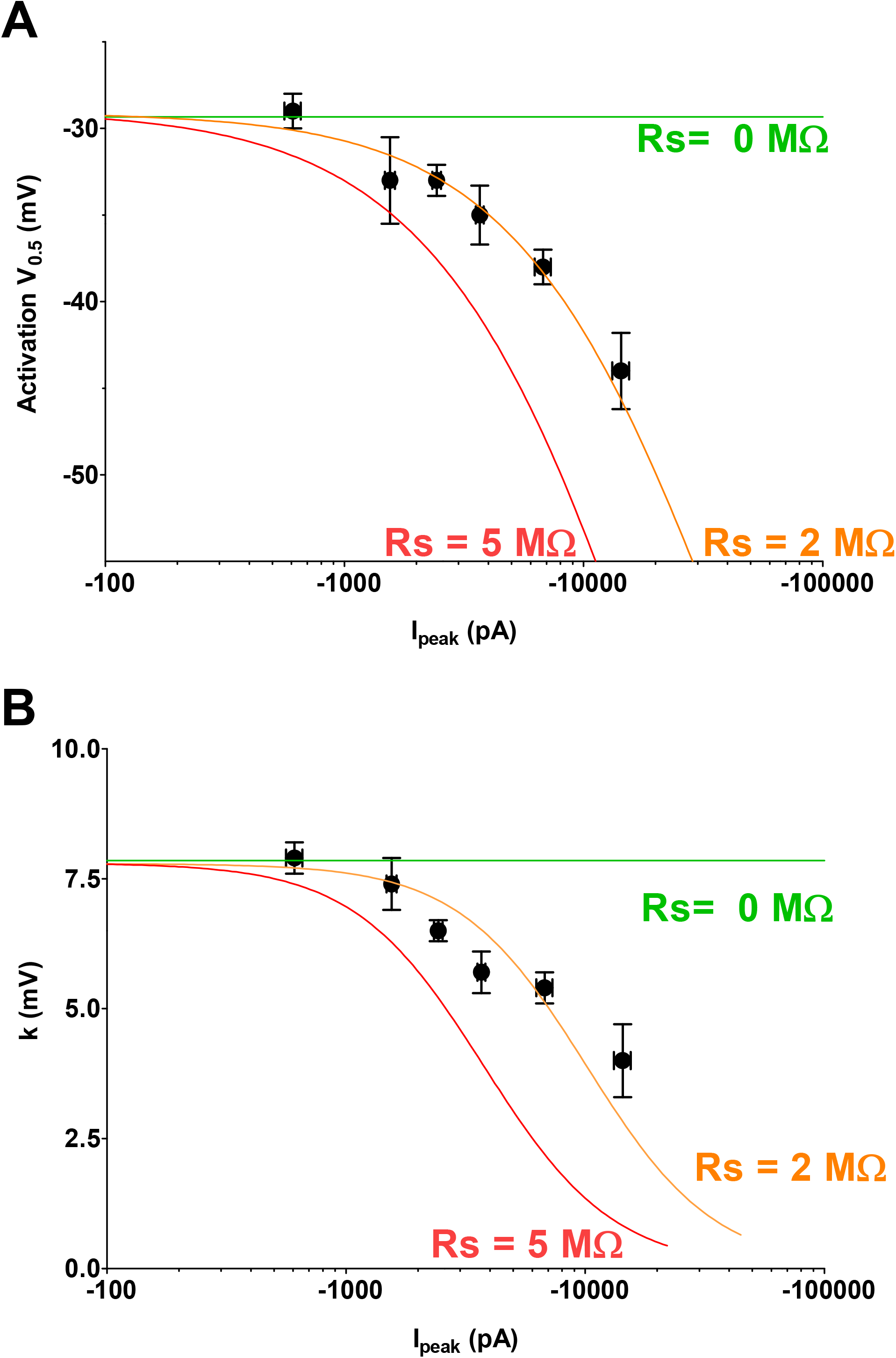
V_0.5_ *vs*. amplitude plots and slope *vs*. amplitude plots of heterologously-expressed Na_v_1.5 currents recorded in COS-7 using manual patch-clamp. **A**. Symbols represent the mean ± sem of pooled experimental values of V_0.5_ as a function of mean ± sem values of current amplitudes. Lines correspond to fit of computed values obtained from the kinetic model that has been optimized in Figure 6 and in which R_S_ has been set to 0, 2 and 5 MΩ (compensated C_m_ = 0.1 pF). **B**. Pooled experimental values of the activation slope as a function of mean ± sem values of current amplitudes.

In order to illustrate how incorrect voltage-clamp due to high series resistance can lead to misinterpretation of the data, we used another set of data of I_Na_ currents obtained from mouse cardiomyocytes in which a “test condition” leads to an increased current amplitude. For the demonstration, we used values of current amplitudes that are frequently published, ranging from −1 to −10 nA, with 80% series resistance compensation. We drew similar plots as in Figure 7, of the experimental activation parameters and we added the model of over-expressed Na_v_1.5 currents generated above. The model does not exactly fit to the data, suggesting that I_Na_ parameters are slightly different in cardiomyocytes and transfected COS-7 cells. Interestingly, however, the exponential fit of the data follows the same trend, parallel to the COS-7 model, suggesting that effects of the treatment on current amplitude is also the cause of the variation of both V_0.5_ and k parameters. This suggests that the differences in V_0.5_ and k parameters between control and test groups may be partly or totally a consequence of the amplitude change (Figure 8).

**Figure 8:**
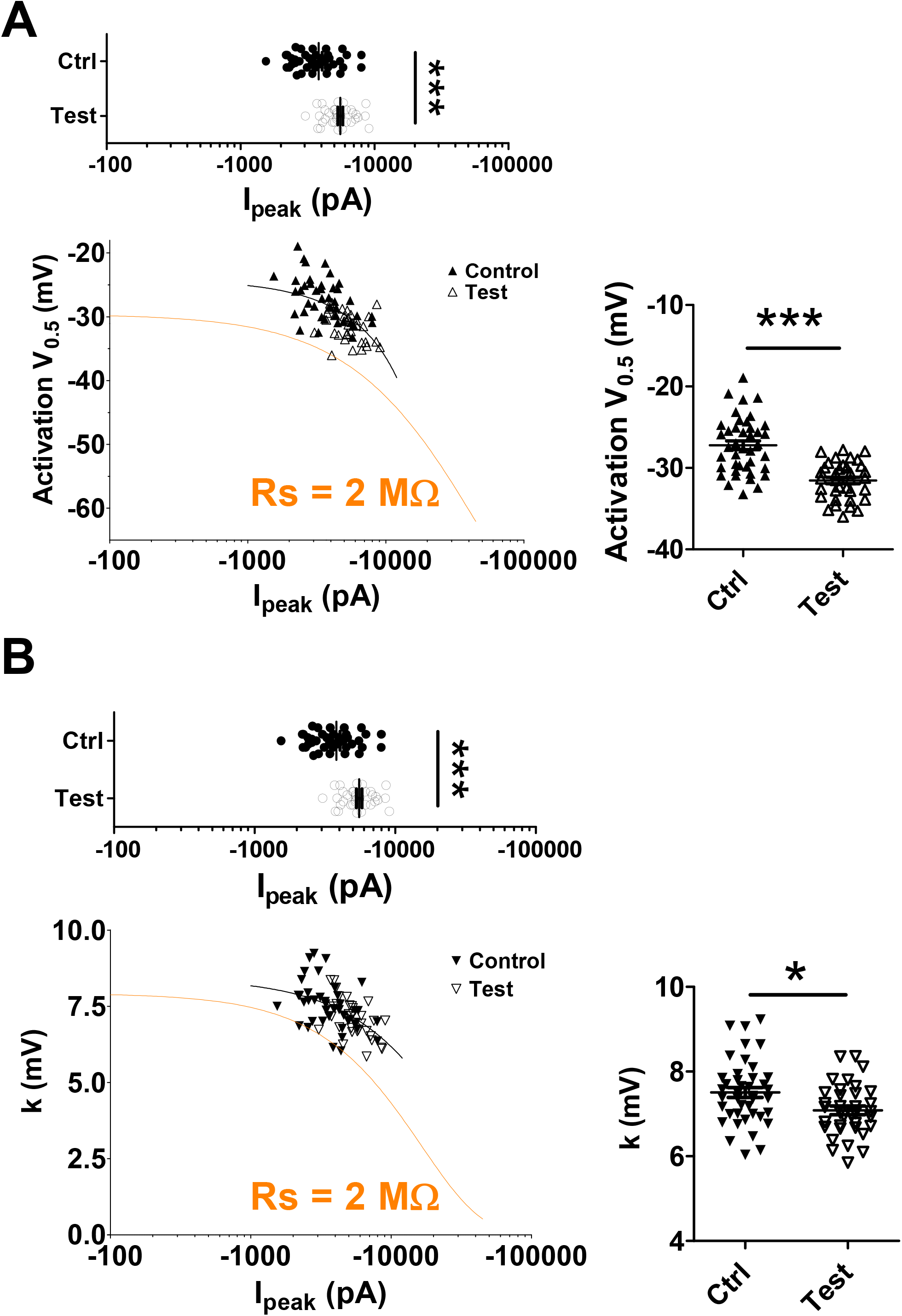
V_0.5_ *vs*. amplitude plots and slope *vs*. amplitude plots of I_Na_ measured from mouse cardiomyocytes using manual patch-clamp. **A**. *Left*, Symbols represent V_0.5_ values as a function of current amplitude values, for control and test conditions. Lines correspond to fit of computed values obtained from the kinetic model that has been optimized for heterologously-expressed Na_v_1.5 currents in COS-7 cells and in which R_S_ has been set to 2 MΩ. Right, Mean ± sem activation V_0.5_ for control and test conditions. ***, p<0.001, student’s t-test. **B**. Same as in A for the slope of the activation curve. *, p<0.05, student’s t-test.

Finally, we used a set of cell data obtained using the automated patch-clamp set-up Syncropatch 384PE, and HEK293 cells stably expressing the Na_v_1.5 channels. The first experimental group has a mean inward current amplitude lower than the reference group of transfected COS-7 cells (−267 ± 67 pA, n = 7 vs −608 ± 48 pA, n = 7, respectively). It has to be noticed that in this amplitude range, activation parameters are more difficult to determine. This is reflected by the large s.e.m. values for mean V_0.5_ and k. We postulate that HEK293 endogenous currents may non-specifically affect the properties of the recorded currents. For the following groups with larger I_Na_ amplitudes, V_0.5_ seems to be stable. Hence, a V_0.5_ value around −25 mV appears to be reliable. When current amplitudes are lower than 3.5 nA, V_0.5_ change is less than 10 mV and k remains greater than 5 mV. Therefore, it is essential to experiment in conditions in which the inward current value is comprised between −300 pA and −3.5 nA when using an automated patch-clamp system, and to exclude data with higher peak current amplitude. These limits are more stringent than for manual patch-clamp as seen above (7 nA), but this is consistent with the limited compensation capabilities of automated patch-clamp systems. Indeed, there is only one amplifier for a large array of cells and each individual R_S_ cannot be measured. Thus, the R_S_ values being different from cell to cell, a series of R_S_ connected in parallel can be partially compensated only but cannot be appropriately compensated by feedback loop without inducing adverse current oscillation.

## DISCUSSION

Even though effects of series resistance have been described very early (Ebihara and Johnson, 1980), a lot of published measured currents are in the range of several nA, which often lead to incorrect voltage-clamp. We developed a simple model, using published kinetic models of ion currents to simulate and describe such a caveat. We used both an inward current generated by a voltage-gated sodium channel and an outward current generated by a voltage-gated potassium channel, both of them characterized by fast activation kinetics. Using these models and experimental recordings, we observed that large series resistance may give erroneous activation curves (Figures 2,3,5,7,9) and dose-response curves (Figure 4). Interpretation of voltage-dependence of activation can further be hindered in cases where treatments of interest alter the current amplitude (Figure 8).

**Figure 9:**
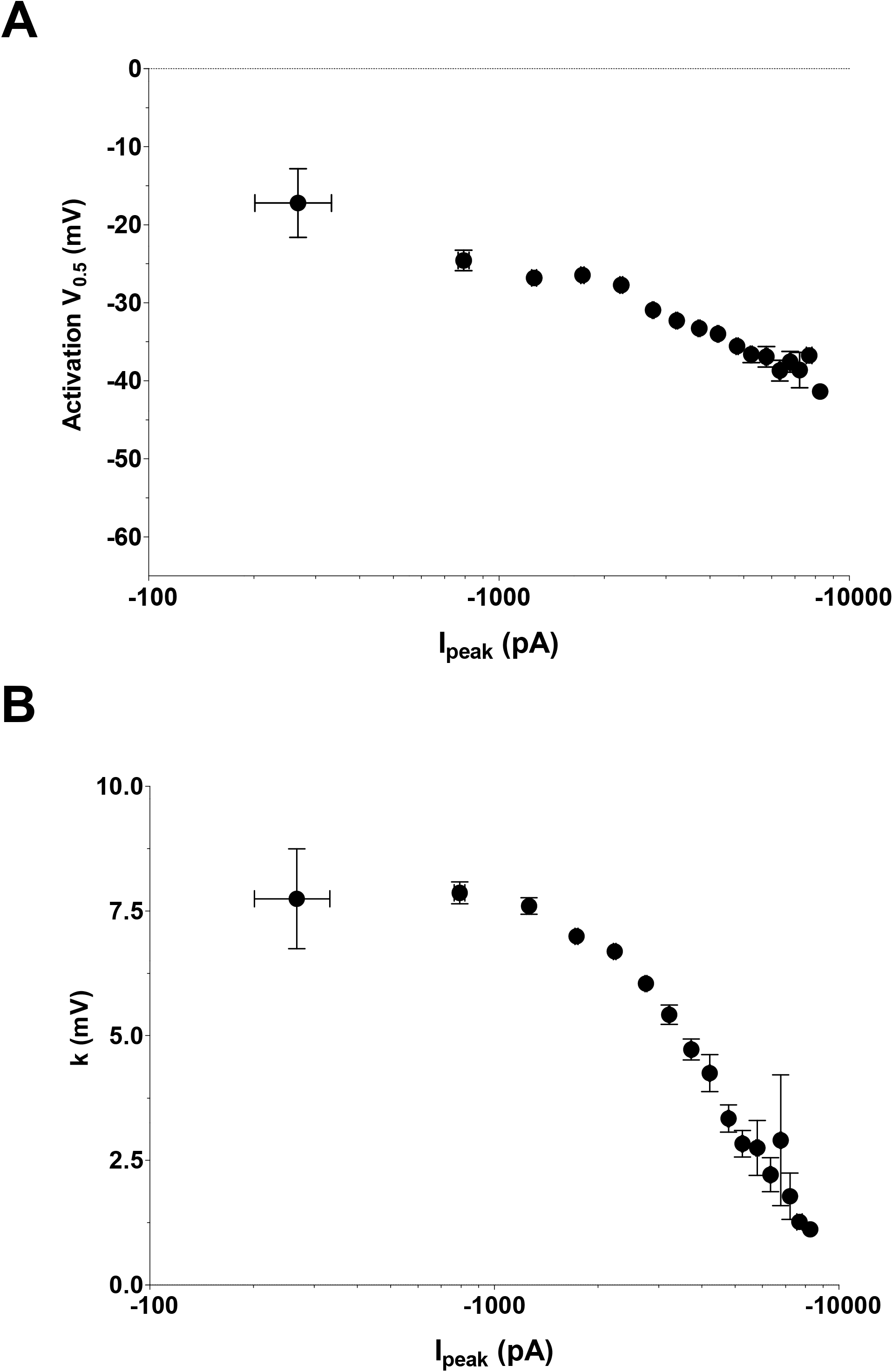
V_0.5_ *vs*. amplitude plots and slope *vs*. amplitude plots for I_Na_ in HEK293 cells stably expressing Na_v_1.5 measured using a Nanion Syncropatch 384PE. **A**. Symbols represent the mean ± sem of pooled values of V_0.5_ as a function of mean ± sem values of current amplitudes. Data points were pooled by 500 pA amplitudes intervals (0-500, 500-1000, etc) **B**. Pooled experimental values of the activation slope as a function of mean ± sem values of current amplitudes.

A similar mathematical model, taking into account the R_S_ impact has been used to study the causes of variability of current recordings obtained from the voltage-gated potassium channel K_v_11.1 (Lei et al., 2019). Here we used such a model to provide a guideline focusing on parameters that the manipulator can easily act on: current amplitude and R_S_.

We observed that the cardiac voltage-gated sodium current I_Na_ is much more sensitive than the cardiac voltage-gated potassium current Ito: the condition with a current amplitude range of 10 nA and a R_S_ of 5 MΩ, shows almost no alteration of the activation curve of the voltage-gated potassium channel (Figure 5B, center and Figure 5C, middle), whereas the same condition in the voltage-gated sodium channel, shows a major alteration of the activation curve (Figure 2B, bottom right and Figure 3, right). The simplest interpretation of this observation may be associated with the fact that, for sodium channels, the increase in sodium entry induced by depolarization further depolarizes the membrane and creates instability. For potassium channels, the increase of potassium outflow induced by depolarization tends to repolarize the membrane and limits instability. However, in extreme cases, repolarization prevents the occurrence of inactivation, leading to delayed inactivation (Figure 5C, bottom right).

For the voltage-gated sodium channel Na_v_1.5, we concluded that it is essential to prevent recording inward current amplitudes greater than 7 nA in which compensated R_S_ is around 2 MΩ to get a reasonable estimate of the activation gate characteristics in manual patch-clamp technique (Figure 7). When using an automated patch system, the limit is lowered to 3.5 nA (Figure 9).

In order to test for activation V_0.5_ changes, induced by drug treatments, mutations, or post-translational modifications which also alter current amplitude, it is advisable to generate plots such as in Figure 8 to determine whether the observed alteration in V_0.5_ and k is a direct consequence of the treatment or an artefactual consequence of the current amplitude change.

To summarize, we suggest simple guidelines for the voltage-gated sodium channel Na_v_1.5:

1. Always compensate R_S_ as much as possible,
2. R_S_ values around 2 MOhm after compensation allows recordings with a maximal current of 7 nA for manual patch-clamp.
3. Using Nanion Syncropatch384PE, recordings with a maximal current of 3.5 nA can be used.

The guidelines are less stringent to record reliable K^+^ outward currents. However, always compensate R_S_ as much as possible. From Figure 5B,C, for R_S_ values up to 5 MΩ after compensation, recordings with a maximal current of 10 nA will be highly reliable. With Nanion Syncropatch384PE, R_S_ values up to 15 MΩ after compensation allow recordings with a maximal current of 10 nA, but inhibitors or various transient transfection conditions should be used to make sure that the measured current amplitude is not saturating due to voltage deviation (Figure 5B, bottom right).

For any current generated by voltage-gated channels, it is mostly judicious to draw activation slope vs. amplitude plots and activation V_0.5_ vs. amplitude plots in a preliminary study to determine adapted conditions, a prerequisite to obtain reliable data and results.

For any other ion-channel type, ligand-, lipid-gated, regulated by second messengers or else, low membrane resistance *i.e*. high expression of active ion channels associated with high R_S_ values will also interfere with adequate voltage command and current measurements.

Several simple adaptations can be made to reach optimal experimental conditions:

1. R_S_ values are much lower when pipettes with low resistance (‘large’ pipettes) are used. When using amplifiers combining R_S_ and C_m_ compensation, suppression of pipette capacitance currents is of high interest since uncompensated pipette capacitance has a detrimental effect on the stability of the series resistance correction circuitry. This can be achieved by the use of borosilicate glass pipettes and wax or Sylgard coating. When using Nanion Syncropatch384PE or other automated patch-clamp systems, low resistance chips are preferred.
2. When over-expressed channels are studied, transfection has to be adapted to produce reasonable amount of channels to generate the desired current amplitude, or, when cell lines stably expressing the channel of interest are used, the clones generating the desired current amplitude range are preferably chosen. Any current, including native currents can be also reduced when pipette and extracellular concentrations of the carried ion are reduced. In addition, the concentration gradient can be changed to limit the electrochemical gradient.

Therefore any patch-clamp experiment needs to be carefully designed to reach appropriate conditions, guaranteeing rigorous analysis of the current.

## METHODS

### Computer models

I_Na_ and I_to_ currents were modeled using a Hodgkin-Huxley model of channel gating based on previously published models (O’Hara et al., 2011).

For cardiac I_Na_, we did not include the slow component of h, which only represents 1% of h inactivation (O’Hara et al., 2011).

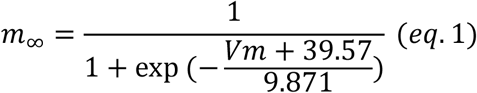

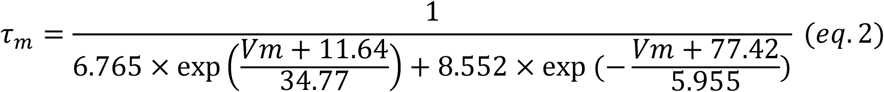

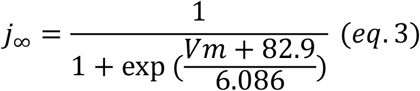

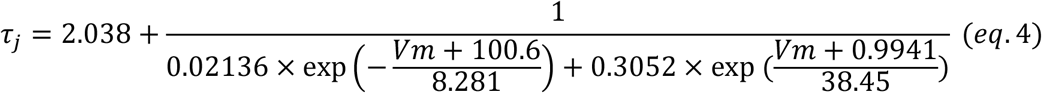

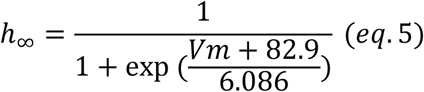

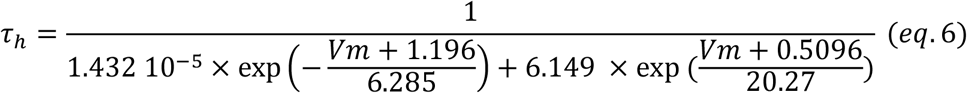

m, h and j were computed as:

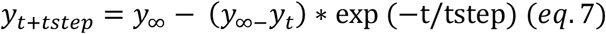

with a time step (tstep) of 0.1 μs

I_Na_ was calculated as follows:

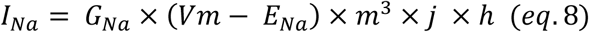

with 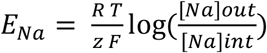, [Na] out = 145 mM and [Na]in = 10 mM

For transfected Na_v_1.5, we adjusted some parameters, shown in bold, to fit the characteristics of the current when peak amplitude is less than 1 nA (cf text and Figure 6).

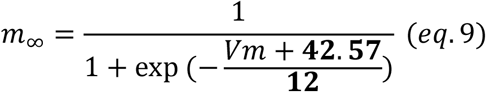

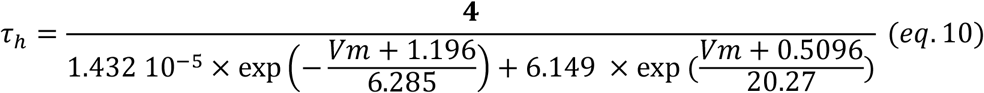

For cardiac I_to_, we did not include the CaMK dependent component, since at low Ca^2+^ pipette concentration (<100 nM), this component is negligible (2%) (O’Hara et al., 2011).

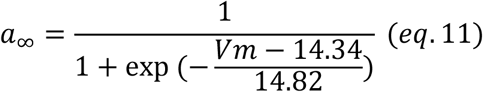

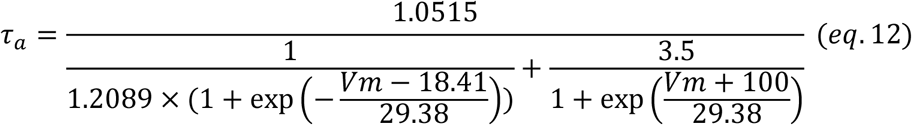

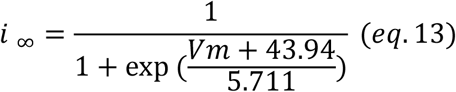

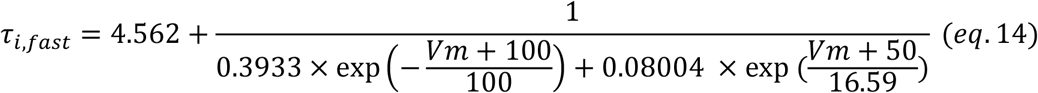

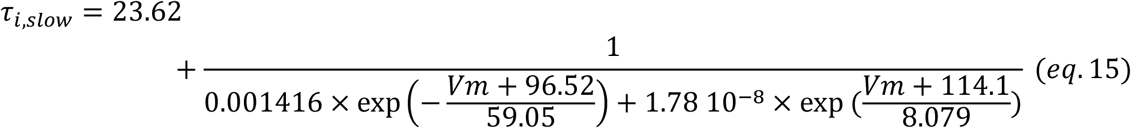

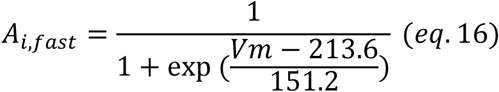

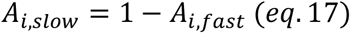

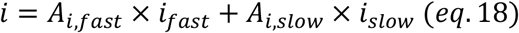

a, *i_fast_* and *i_low_* were computed as:

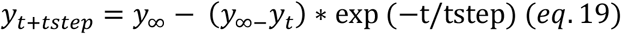

with a time step (tstep) of 0.1 μs

I_to_ was calculated as follows:

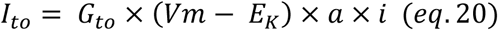

with 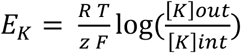, [K]out = 5 mM and [K]in = 145 mM

For details on the kinetic models, please see (O’Hara et al., 2011)

Voltage error caused by the presence of R_S_ was computed as follows at each time step:

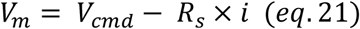

We hypothesized that the amplifier response time was not limiting. Membrane capacitance used was 25 pF and considered electronically compensated. Space clamp was considered negligible in small cells like COS-7 cells but it is worth mentioning that it should be taken into account in bigger cells such as cardiomyocytes and in cells with complex morphologies such as neurons.

### Cell culture and transfection

The African green monkey kidney-derived cell line, COS-7, was obtained from the American Type Culture Collection (CRL-1651) and cultured in Dulbecco’s modified Eagle’s medium (GIBCO) supplemented with 10% fetal calf serum and antibiotics (100 IU/mL penicillin and 100 μg/mL streptomycin) at 5% CO_2_ and 95% air, maintained at 37°C in a humidified incubator. Cells were transfected in 35-mm Petri dishes when the culture reached 50-60% confluence, with DNA (2 μg total DNA) complexed with jetPEI (Polyplus transfection) according to the standard protocol recommended by the manufacturer. COS-7 cells were co-transfected with 200 ng of *pCI-SCN5A* (NM_000335.4), 200 ng of pRC-*SCN1B* (NM_001037) (kind gifts of AL George, Northwestern University, Feinberg School of Medicine) and 1.6 μg pEGFP-N3 plasmid (Clontech). Cells were re-plated onto 35-mm Petri dishes the day after transfection for patch-clamp experiments. HEK293 cells stably expressing hNa_v_1.5 were cultured in Dulbecco’s Modified Eagle’s Medium (DMEM) supplemented with 10% fetal calf serum, 1 mM pyruvic acid, 4 mM glutamine, 400 μg/ml of G418 (Sigma), 10 U/mL penicillin and 10 μg/mL streptomycin (Gibco, Grand Island, NY), and incubated at 37°C in a 5% CO_2_ atmosphere.

### Statement on the use of mice

All investigations conformed to directive 2010/63/EU of the European Parliament, to the Guide for the Care and Use of Laboratory Animals published by the US National Institutes of Health (NIH Publication No. 85-23, revised 1985) and to local institutional guidelines.

### Cardiomyocyte preparation

Single cardiomyocytes were isolated from the ventricles of adult (8-14 weeks) mice by enzymatic dissociation and mechanical dispersion according to a modified procedure of established methods. Briefly, mice were injected with heparin (5000 units/kg body weight) 30 minutes before sacrifice by cervical dislocation. Hearts were excised, and perfused retrogradely through the aorta with a solution containing (in mM): NaCl, 113; KCl, 4.7; MgSO_4_, 1.2; KH_2_PO_4_, 0.6; NaH_2_PO_4_, 0.6; HEPES, 10; NaHCO_3_, 1.6; taurine, 30; glucose, 20 (pH 7.4 with NaOH). Hearts were subsequently digested for 11 minutes with the same solution supplemented with 0.08 mg/mL Liberase TM Research Grade (Sigma). Following digestion, the perfusion was stopped, the atria were removed, and the ventricles were dispersed by gentle trituration. Cell suspensions were filtered to remove large undissociated tissue fragments, and resuspended in solutions containing 10 mg/mL bovine serum albumin and Ca^2+^ concentrations successively increasing from nominally 0 to 0.2, 0.5 and 1 mM. Cells were then resuspended in medium-199 (Sigma) supplemented with 5% foetal bovine serum, 10 mM 2,3-Butanedione monoxime, 100 U/ml penicillin and 100 μg/ml streptomycin, plated on laminin-coated dishes, and incubated in 37°C, 5% CO_2_: 95% air incubator. After 1-hour plating, culture medium was replaced by medium-199 supplemented with 0.1% bovine serum albumin, 10 mM 2,3-Butanedione monoxime, 1X Insulin/Transferrin/Sodium Selenite (Sigma), 1X Chemically Defined Lipid Concentrate (Gibco), 0.5 μM cytochalasine D (Sigma), 100 U/mL penicillin and 100 μg/mL streptomycin until electrophysiological experiments were performed.

### Manual electrophysiology on transfected COS-7 cells

One or 2 days after splitting, COS-7 cells were mounted on the stage of an inverted microscope and constantly perfused by a Tyrode solution maintained at 22.0 ± 2.0°C at a rate of 1-3 mL/min; HEPES-buffered Tyrode solution contained (in mmol/L): NaCl 145, KCl 4, MgCl_2_ 1, CaCl_2_ 1, HEPES 5, glucose 5, pH adjusted to 7.4 with NaOH. During Na^+^ current recording, the studied cell was locally superfused with extracellular medium containing (in mmol/L): NaCl, 145; CsCl, 4; CaCl_2_, 1; MgCl_2_, 1; HEPES, 5; glucose, 5; pH adjusted to 7.4 with NaOH. Patch pipettes (tip resistance: 0.8 to 1.3 MΩ) were pulled from soda lime glass capillaries (Kimble-Chase) and coated with dental wax to decrease pipette capacitive currents. The pipette was filled with Na^+^ intracellular medium containing (in mmol/L): CsCl, 80; gluconic acid, 45; NaCl, 10; MgCl_2_, 1; CaCl_2_, 2.5; EGTA, 5; HEPES, 10; pH adjusted to 7.2 with CsOH. Stimulation and data recording were performed with pClamp 10, an A/D converter (Digidata 1440A) and an Axopatch 200B (all Molecular Devices) or an Alembic amplifier (Alembic Instruments). Currents were acquired in the whole-cell configuration, filtered at 10 kHz and recorded at a sampling rate of 33 kHz. Before Series resistance compensation, a series of 50 25-ms steps were applied from −70 mV to −80 mV to subsequently off-line calculate more precise C_m_ and R_S_ values from the recorded current. To generate the Na_v_1.5 activation curve, the membrane was depolarized from a holding potential of −100 mV to values between −80 mV and +50 mV (+5-mV increment) for 50 ms, every 2 s. Activation curves were fitted by a Boltzmann equation: G = G_max_ / (1 + exp (− (V_m_ − V_0.5_) / k)), in which G is the conductance, V_0.5_ is the membrane potential of half-activation and k is the slope factor.

### Electrophysiology on cardiomyocytes

Whole-cell Na_v_ currents were recorded at room temperature 48 hours after cell isolation using an Axopatch 200B amplifier (Axon Instruments, Molecular Devices, San Jose, CA). Voltage-clamp protocols were applied using the pClamp 10.2 software package interfaced to the electrophysiological equipment using a Digidata 1440A digitizer (all Axon Instruments). Current signals were filtered at 10 kHz prior to digitization at 50 kHz and storage. Patch-clamp pipettes were fabricated from borosilicate glass (OD: 1.5 mm, ID: 0.86 mm, Sutter Instrument, Novato, CA) using a P-97 micropipette puller (Sutter Instrument), coated with wax, and fire-polished to a resistance between 0.8 and 1.5 MΩ when filled with internal solution. The internal solution contained (in mM): NaCl 5, CsF 115, CsCl 20, HEPES 10, EGTA 10 (pH 7.35 with CsOH, ~300 mosM). The external solution contained (in mM): NaCl 10, CsCl 5, N-Methyl-D-glucamine-Cl 104, TEA-Cl (tetraethylammonium chloride) 25, HEPES 10, Glucose 5, CaCl_2_ 1, MgCl_2_ 2 (pH 7.4 with HCl, ~300 mosM). All chemicals were purchased from Sigma. After establishing the whole-cell configuration, ten minutes were allowed to allow stabilization of voltage-dependence of activation and inactivation properties. During this time, 25-ms voltage steps to ± 10 mV from a holding potential (HP) of −70 mV were applied to allow measurement of whole-cell membrane capacitances, input and series resistances. After compensation of series resistance (80%), the membrane was held at a HP of −120 mV, and the voltage-clamp protocol was carried out as indicated below.

Data were compiled and analyzed using ClampFit 10.2 (Axon Instruments), Microsoft Excel, and Prism (GraphPad Software, San Diego, CA). Whole-cell membrane capacitances (Cm) were determined by analyzing the decays of capacitive transients elicited by brief (25 ms) voltage steps to ± 10 mV from the HP (−70 mV). Input resistances were calculated from the steady-state currents elicited by the same ±10 mV steps (from the HP). Series resistances were calculated by dividing the decay time constants of the capacitive transients (fitted with single exponentials) by the Cm. To determine peak Na^+^ current-voltage relationships, currents were elicited by 50-ms depolarizing pulses to potentials ranging from −80 to +40 mV (presented at 5-s intervals in 5-mV increments) from a HP of −120 mV. Peak current amplitudes were defined as the maximal currents evoked at each voltage, and were subsequently leak-corrected. To analyze voltage-dependence current activation properties, conductances (G) were calculated, and conductance-voltage relationships were fitted with a Boltzmann equation

### High-throughput electrophysiology

Automated patch-clamp recordings were performed using the SyncroPatch 384PE from Nanion (München, Germany). Single-hole, 384-well recording chips with medium resistance (4.77 ± 0.01 MΩ, n=384) were used for recordings of HEK293 cells stably expressing human Na_v_1.5 channel (300 000/mL) in whole-cell configuration. Pulse generation and data collection were performed with the PatchControl384 v1.5.2 software (Nanion) and the Biomek v1.0 interface (Beckman Coulter). Whole-cell recordings were conducted according to the recommended procedures of Nanion. Cells were stored in a cell hotel reservoir at 10°C with shaking speed at 60 RPM. After initiating the experiment, cell catching, sealing, whole-cell formation, buffer exchanges, recording, and data acquisition were all performed sequentially and automatically. The intracellular solution contained (in mM): 10 CsCl, 110 CsF, 10 NaCl, 10 EGTA and 10 HEPES (pH 7.2, osmolarity 280 mOsm), and the extracellular solution contained (in mM): 60 NaCl, 4 KCl, 100 NMDG, 2 CaCl2, 1 MgCl2, 5 Glucose and 10 HEPES (pH 7.4, osmolarity 298 mOsm). Whole-cell experiments were performed at a holding potential of −100 mV at room temperature (18-22°C). Currents were sampled at 20 kHz. Activation curves were built by 50 ms-lasting depolarization from −80 mV to 70 mV (+5 mV increment), every 5 s. Activation curves were fitted by Boltzmann equation. Stringent criteria were used to include individual cell recordings for data analysis (seal resistance > 0.5 GΩ and estimated series resistance < 10 MΩ).

## ACKNOWLEDGEMENTS

We are indebted to Dr. Massimo Mantegazza for providing the stable cell line expressing human Na_v_1.5. The authors wish to thank Drs Massimo Mantegazza and Flavien Charpentier for their critical reading of the manuscript. M. De Waard thanks the Agence Nationale de la Recherche for its financial support to the laboratory of excellence “Ion Channels, Science and Therapeutics” (grant N° ANR-11-LABX-0015). This work was supported by the Fondation Leducq in the frame of its program of ERPT equipment support (purchase of an automated patch-clamp system), by a grant “New Team” of the Région Pays de la Loire to M. De Waard, and by a European FEDER grant in support of the automated patch-clamp system of Nanion. The salary of S. Nicolas is supported by the Fondation Leducq, while the fellowship of J. Montnach is provided by a National Research Agency Grant to M. De Waard entitled OptChemCom (grant N° ANR-18-CE19-0024-01). C. Marionneau thanks the Agence Nationale de la Recherche [ANR-15-CE14-0006-01 and ANR-16-CE92-0013-01]. M. Lorenzini was supported by a Groupe de Réflexion sur la Recherche Cardiovasculaire-Société Française de Cardiologie predoctoral fellowship [SFC/GRRC2018].

## AUTHOR CONTRIBUTION

Jérome Montnach and Sébastien Nicolas carried out the automated patch-clamp experiments on Na_v_1.5, under the supervision of Michel De Waard. Maxime Lorenzini and Adrien Lesage carried out the patch-clamp experiments on mouse cardiomyocytes, under the supervision of Céline Marionneau. Isabelle Simon and Eléonore Moreau carried out the manual patch-clamp experiments on transfected COS-7 cells, under the supervision of Isabelle Baró. Gildas Loussouarn generated the model and wrote the manuscript.

## References

Ebihara L, Johnson EA. 1980. Fast sodium current in cardiac muscle. A quantitative description. Biophysical Journal 32:779–790. doi:10.1016/S0006-3495(80)85016-8

Hille B. 2001. Ion Channels of Excitable Membranes, 3rd edition. ed. Sinauer Associates (distributed by W.H. Freeman).

Lei CL, Clerx M, Whittaker DG, Gavaghan DJ, de Boer TP, Mirams GR. 2019. Accounting for variability in ion current recordings using a mathematical model of artefacts in voltage-clamp experiments (preprint). Biophysics. doi:10.1101/2019.12.20.884353

O’Hara T, Virág L, Varró A, Rudy Y. 2011. Simulation of the Undiseased Human Cardiac Ventricular Action Potential: Model Formulation and Experimental Validation. PLoS Comput Biol 7:e1002061. doi:10.1371/journal.pcbi.1002061

Oliver KL, Franceschetti S, Milligan CJ, Muona M, Mandelstam SA, Canafoglia L, Boguszewska-Chachulska AM, Korczyn AD, Bisulli F, Di Bonaventura C, Ragona F, Michelucci R, Ben-Zeev B, Straussberg R, Panzica F, Massano J, Friedman D, Crespel A, Engelsen BA, Andermann F, Andermann E, Spodar K, Lasek-Bal A, Riguzzi P, Pasini E, Tinuper P, Licchetta L, Gardella E, Lindenau M, Wulf A, Møller RS, Benninger F, Afawi Z, Rubboli G, Reid CA, Maljevic S, Lerche H, Lehesjoki A-E, Petrou S, Berkovic SF. 2017. Myoclonus epilepsy and ataxia due to *KCNC 1* mutation: Analysis of 20 cases and K + channel properties. Ann Neurol 81:677–689. doi:10.1002/ana.24929

Ranjan R, Logette E, Marani M, Herzog M, Tâche V, Scantamburlo E, Buchillier V, Markram H. 2019. A Kinetic Map of the Homomeric Voltage-Gated Potassium Channel (Kv) Family. Front Cell Neurosci 13:358. doi:10.3389/fncel.2019.00358

Sherman AJ, Shrier A, Cooper E. 1999. Series Resistance Compensation for Whole-Cell Patch-Clamp Studies Using a Membrane State Estimator. Biophysical Journal 77:2590–2601. doi:10.1016/S0006-3495(99)77093-1

Wang GK, Russell G, Wang S-Y. 2013. Persistent human cardiac Na ^+^ currents in stably transfected mammalian cells: Robust expression and distinct open-channel selectivity among Class 1 antiarrhythmics. Channels 7:263–274. doi:10.4161/chan.25056

