## Supplemental Data for "Computer modeling of whole-cell voltage-clamp analyses to delineate guidelines for good practice of manual and automated patch-clamp"

**Relationship between ion current and the quality of voltage-clamp summarized in a limited number of equations.**

For a given ionic species, the current *I* depends on the driving force applied to the carried ion to cross the channel. For $Na^+$ current:

$$I_{Na} = \frac{(V_m - E_{Na})}{R_m} \qquad \text{eq. s1}$$

with $V_m$ the voltage difference across the cell membrane; $R_m$, the membrane resistance for $Na^+$ (inversely proportional to the number of $Na^+$ channels in the open state); $E_{Na}$, the reversal potential determined by the Nernst equation:

$$E_{Na} = \frac{RT}{zF} \ln([Na^+]_{out}/[Na^+]_{in}) \qquad \text{eq. s2}$$

where $R$ is the universal gas constant (8.314 Joules per Kelvin per mole); *T*, the temperature in Kelvin; *z*, the valence of the ionic species (+1 for $Na^+$); *F*, the Faraday constant (96485 Coulombs per mole). $[Na^+]_{out}$ is the concentration of $Na^+$ in the extracellular solution and $[Na^+]_{in}$ in the intracellular medium (pipette solution).

The series resistance is a sum of resistances dependent on the pipette tip size and access into the cell. According to the Ohm law: $V = R \times I$

For $R_S$ and $R_m$, two resistances in series:

$V_{cmd} = V_S + V_m$ and, from eq. s1, $V_m = R_m \times I_{Na} + E_{Na}$

$$V_{cmd} = (R_S \times I_{Na}) + (R_m \times I_{Na}) + E_{Na}$$

If most of the channels are closed or lowly expressed, $R_m \gg R_S$ and $V_S$ is negligible when compared to $V_m$. In this case, $V_m \equiv V_{cmd}$.

When channels are opening, $R_m$ decreases and *I* increases. As long as $R_S$ remains low by using pipettes with large tip, and $R_m$ remains several log higher than $R_S$ (by maintaining a low channel expression), $V_m$ remains fairly close to $V_{cmd}$. But if the channels are highly expressed, $R_m$ can reach values in the range of $R_S$. In these conditions, $V_S$ cannot be neglected when compared to $V_m$ and $V_m$ becomes a fraction of $V_{cmd}$, only.

For native $Na^+$ channels in cardiomyocytes, for example, the current amplitude can be lowered by reducing $E_{Na}$ (eq. s2) *i.e.* by decreasing the concentration gradient, and by reducing the number of ions crossing the channels by partly substituting them with an equivalent non-permeant ion in the solutions (here $Cs^+$ for $Na^+$).

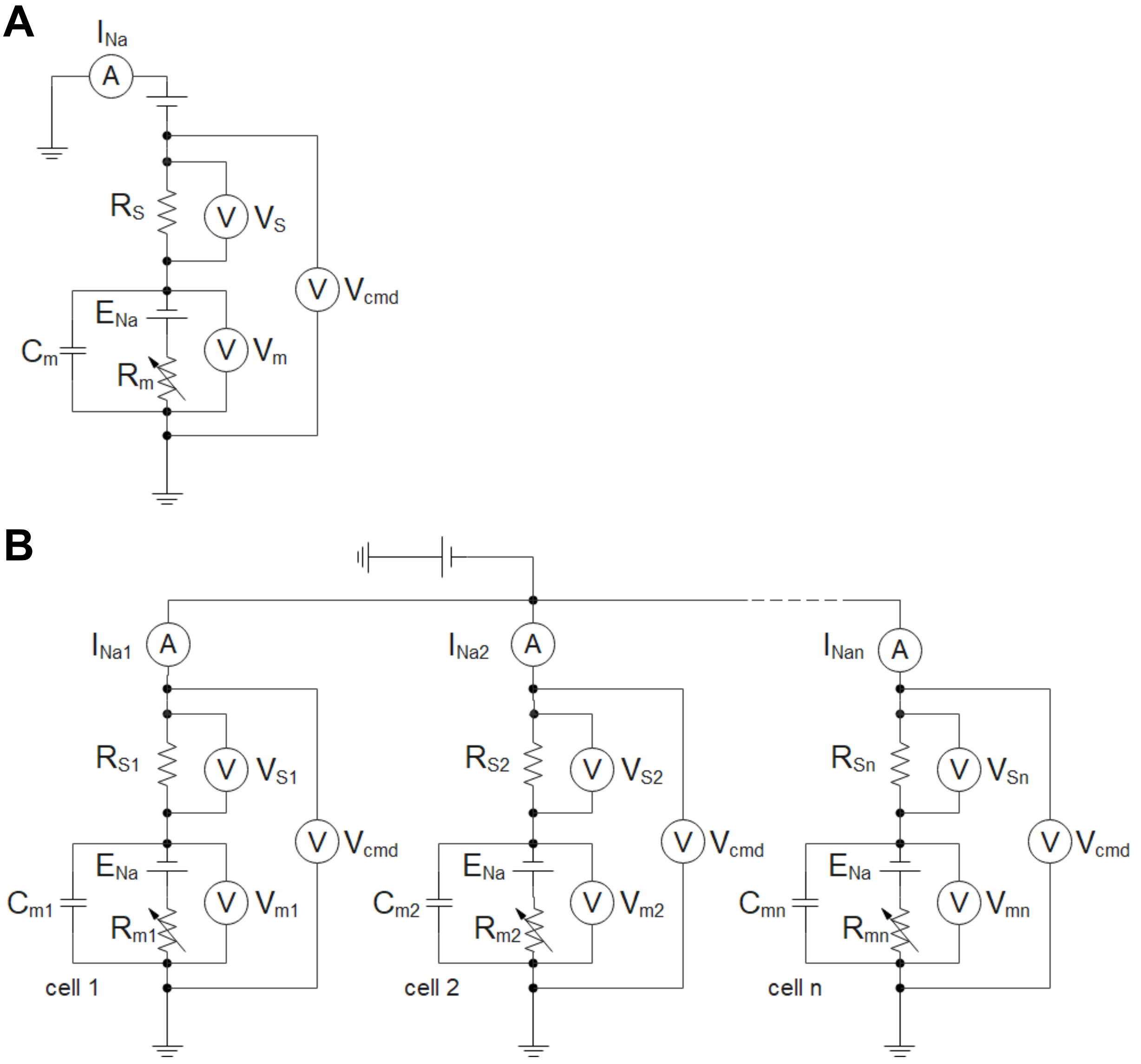


**Figure S1: Schemes of the electrical circuits in voltage-clamp to measure the $Na^+$ current ($I_{Na}$),** illustrating how the potential clamped between the two electrodes ($V_{cmd}$) is split between the membrane potential ($V_m$) and a potential drop generated at the series resistance ($R_S$) for a cell. **A.** Manual patch-clamp. Pipette capacitance and leak current due to imperfect seal between the pipette and the cell have been omitted for the sake of clarity. $R_m$: cell resistance for $Na^+$. This value depends on the $Na^+$ channel expression level and the fraction of expressed $Na^+$ channels that are open (higher is the value, less are the active $Na^+$ channels). The fraction of channel in the open state depends on voltage and time. $E_{Na}$: value of the reversal potential for $Na^+$ (determined by intra- and extracellular $Na^+$ concentrations according to the Nernst equation); Cm: cell capacitance (higher is the value, larger is the cell). B. Automated patch-clamp. The circuit of only one amplifier is shown. One amplifier is used per n cells. As in A, leak current due to imperfect seal between the chip and the cell has been omitted for the sake of clarity.
